# VEGFA’s distal enhancer regulates its alternative splicing in CML

**DOI:** 10.1101/2021.01.09.426072

**Authors:** Sara Dahan, Klil Cohen, Mercedes Bentata, Eden Engal, Ahmad Siam, Gillian Kay, Yotam Drier, Shlomo Elias, Maayan Salton

## Abstract

Enhancer demethylation in leukemia and lymphoma was shown to lead to overexpression of genes which promote cancer characteristics. The vascular endothelial growth factor A (VEGFA) enhancer, located 157 Kb downstream of its promoter, is demethylated in chronic myeloid leukemia (CML). VEGFA has several alternative splicing isoforms with different roles in vascularization and cancer progression. Since transcription and splicing are coupled in space and time, we hypothesized that the VEGFA enhancer can also regulate the gene’s alternative splicing to contribute to the pathology of CML. Our results show that mutating the VEGFA +157 enhancer promotes exclusion of exons 6b and 7 and activating the enhancer by tethering a chromatin activator has the opposite effect. In line with these results, CML patients display high enhancer activity and inclusion of VEGFA exons 6b and 7. To search for a key protein connecting transcription with alternative splicing, we analyzed 161 chromatin immunoprecipitation (ChIP)-seq experiments for DNA binding proteins and found that PML and CCNT2 bind VEGFA’s promoter and enhancer. CCNT2 is a positive regulator of RNA polymerase II (RNAPII) transcription elongation and indeed its silencing promotes exclusion of exons 6b and 7. Slowing down RNAPII elongation promotes exclusion of exons 6b and 7. Thus our results suggest that VEGFA’s +157 enhancer regulates its alternative splicing by increasing RNAPII elongation rate via CCNT2. Our work demonstrates the importance of the interplay between transcription and alternative splicing.

## Introduction

Transcriptional enhancers are major regulators of tissue-specific gene expression. During cellular differentiation, enhancers that control the expression of genes involved in lineage specification become active, while others are deactivated. Thus, the maintenance of cell-type-specific enhancer activation is critical to prevent disease. The disruption of enhancer activity, through genetic or epigenetic alterations, can impact cell-type-specific functions, and promote tumorigenesis of various cancers [1].

Enhancer DNA methylation was shown to play an important part in maintaining and modifying enhancer activity during differentiation. Vascular endothelial growth factor A (VEGFA) enhancer, located 157 Kb downstream to the gene’s promoter (+157), is active in embryonic stem cells, but during hematopoiesis it becomes methylated and inactive [2]. In chronic myeloid leukemia (CML), the enhancer is demethylated and VEGFA is overexpressed [2]. VEGFA over-expression has been shown to be correlated with a range of tumor types [3-5] and to promote growth and survival of vascular endothelial cells. While vascularization is critical in solid tumors to allow for oxygen and nutrient flux [6], it has been shown to be present in hematological malignancies as well [7, 8]. Interestingly, VEGFA alternative splicing changes in many types of cancer and alternative splicing in general is known to be affected by expression levels.

CML is triggered by a Philadelphia chromosome encoding the BCR-ABL oncogenic fusion protein with constitutive and aberrant tyrosine kinase activity [9]. The introduction of tyrosine kinase inhibitors (TKIs) revolutionized the management of CML and patients’ life expectancy is now approaching normality [10]. Regardless, some CML patients do not respond to TKIs and progress to acute leukemia [11]. BCR-ABL was shown to induce the expression of hypoxia-inducible factor-1 (HIF-1), a transcription factor binding to the VEGFA promoter and enhancer regions [12, 13]. In addition, drugs targeting BCR-ABL reduce VEGFA expression [14, 15]. In turn VEGFA overexpression is associated with the clinical characteristics of CML [16]. High levels of VEGFA were recorded in the blood as well as the bone marrow of CML patients and were correlated with high vascularization and proliferation leading to the pathology of CML [17]. VEGFA alternative splicing isoforms have different roles in vascularization and cancer progression, but their expression in CML has not been studied.

Alternative splicing produces proteins with different functions and therefore is tightly regulated. Changes to alternative splicing are common in many diseases, including cancer; alternative splicing misregulation in cancer cells promotes the formation of cancer-driving isoforms [18]. These isoforms have been shown to be involved in each of the hallmarks of cancer [reviewed in 19]; and splicing inhibitors can serve as a therapy for many types of cancers [20, 21].

A strong connection between transcription and splicing has been known for 30 years [22]. Manipulating a gene’s promoter or positioning a synthetic enhancer next to a gene can alter alternative splicing [22-24]. In addition, slow kinetics of RNAPII elongation promotes exon inclusion; the included exon is the first to transcribe and consequently the first to splice [25]. On the contrary, RNAPII fast kinetics promotes exon exclusion as it exposes additional splice sites [25]. The opposite effect can also be observed whereby slow RNAPII kinetics promotes exon exclusion. This effect was attributed to an inhibitory splicing factor that gains a binding opportunity when RNAPII is slow to transcribe [26]. This connection between transcription and alternative splicing suggests that a change in transcription in cancer cells can lead to a change in gene expression as well as alternative splicing.

Our results demonstrate that alternative splicing of VEGFA is co-regulated with its expression by its +157 enhancer. VEGFA’s major splicing isoforms are VEGFA_121_, VEGFA_165_ and VEGFA_189_; their relative abundance in cancer is correlated with pathogenesis but their functional roles are poorly characterized [27-30] (**Fig. 1A**). To learn about the connection of the VEGFA enhancer to its alternative splicing, we used the CML cell line, K562, mutated at the VEGFA enhancer using the CRISPR/Cas9 genome-editing system. It has previously been shown that mutations at this site reduce the total mRNA amount of VEGFA [2] and we further found that it regulates its alternative splicing to promote exclusion of VEGFA exons 6b and 7, which translates to VEGFA_121_. Activating the enhancer using CRISPR/dCas9 to tether p300 gave the opposite result. In line with this result, CML patients show high expression of +157 eRNA and inclusion of VEGFA exons 6b and 7 which translates to VEGFA_189_. To study the mechanism of enhancer regulation of alternative splicing, we altered RNAPII transcription elongation kinetics and monitored alternative splicing. We found that similar to mutations at the +157 enhancer, slowing down RNAPII elongation promotes exclusion of VEGFA exons 6b and 7. Finally we identified PML and CCNT2 as chromatin factors binding to VEGFA promoter and +157 enhancer and as regulators of its alternative splicing. Since CCNT2 is a known positive regulator of RNAPII, our results suggest that CCNT2 mediates the RNAPII elongation kinetics caused by the +157 enhancer, which regulates VEGFA alternative splicing. Ultimately our work identifies for the first time a connection between an endogenous enhancer and alternative splicing regulation of its target gene.

**Fig 1:**
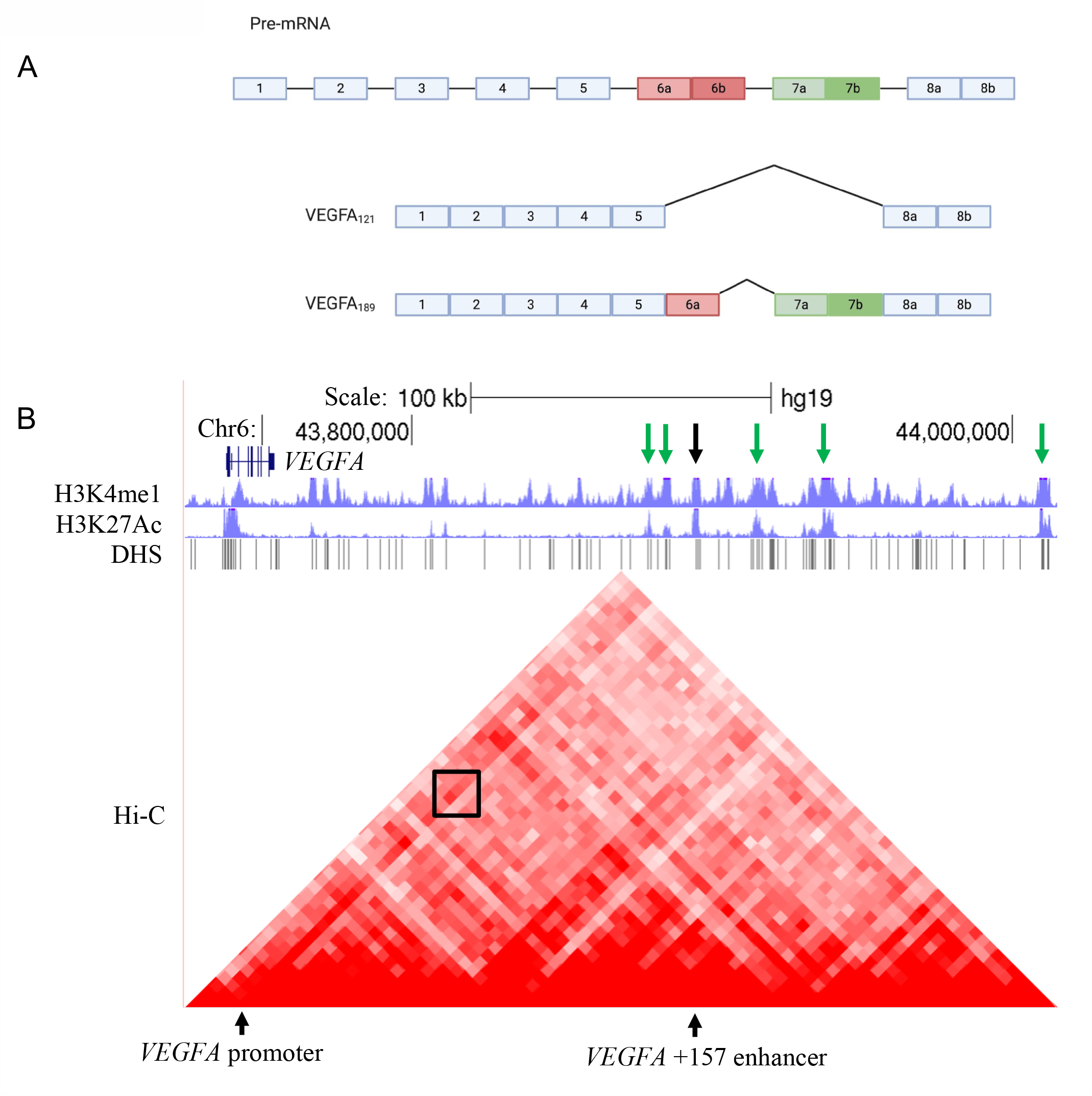
VEGFA isoforms and +157 enhancer region. **A**. Schematic representation of the VEGFA gene and two of its known isoforms. Rectangles: exons, black lines: introns. **B**. ChIP-seq tracks for H3K27ac and H3K4me1 including DHS and Hi-C data at the VEGFA locus in K562 cells. VEGFA +157 enhancer is depicted with a black arrow, and enriched for H3K27ac and H3K4me1. The Hi-C data suggest a sub-TAD between VEGFA +157 enhancer and the promoter, marked with a black box. Green arrows depict additional enhancers with sub-TADs to VEGFA promoter.

## Results

### VEGFA +157 enhancer is marked by H3K4me1, H3K27ac and DNase-seq and loops to its promoter

Enhancers are marked with specific histone modifications, including monomethylation of histone H3 on Lys 4 (H3K4me1) and acetylation of histone H3 on Lys 27 (H3K27ac). They are also associated with regions of nucleosome depletion, exhibiting high sensitivity to DNA nucleases such as the DNase I, forming DNase hypersensitivity sites (DHS). We analyzed histone modification and DHS data in K562 cells (ENCODE Project Consortium [31]) and found that indeed the *VEGFA* +157 enhancer is marked by H3K4me1 and H3K27ac and exhibits multiple DHS sites. Moreover, we analyzed K562 Hi-C chromosome conformation capture data [31] and identified a stripe architecture [32] at the VEGFA promoter, with marked interaction between the promoter and the *VEGFA* +157 enhancer, suggesting they frequently interact (**Fig. 1B and Supplementary Fig. S1**). In addition, we identified five additional putative VEGFA enhancers with similar characteristics (**Fig. 1B**).

### Mutations in VEGFA +157 enhancer promote exclusion of VEGFA exons 6b &7

To check whether VEGFA’s enhancer regulates its alternative splicing, we used the CML cell line, K562, with mutations in VEGFA + 157 enhancer [2]. We used two clones, 9 and 26, generated using the CRISPR/Cas9 genome-editing system. Clone 9 has a single nucleotide insertion at the three chromosome 6 copies in this cell line, while clone 26 has various deletions (**Supplementary Fig. S1**). We began by measuring VEGFA total mRNA in the parental K562 cell line and in clones 9 and 26. As was described before [2], we detected a 20-30% reduction in VEGFA total mRNA following the impairment of the +157 enhancer (**Fig. 2A**). To check the transcription level of VEGFA in these cells, we used primers spanning the exon-intron junction to measure nascent pre-mRNA. The level of transcription had similar trend to VEGFA total mRNA amount indicating lower transcription rate and not a change in mature mRNA half-life (**Supplementary Fig. S2A)**. Furthermore, we detected exclusion of VEGFA exons 6b and 7 in mutated +157 enhancer cells (**Fig. 2B**). Our results suggest that VEGFA +157 enhancer promotes inclusion of exons 6b and 7 giving rise to VEGFA_121_ and a reduction in VEGFA_189_.

**Fig 2:**
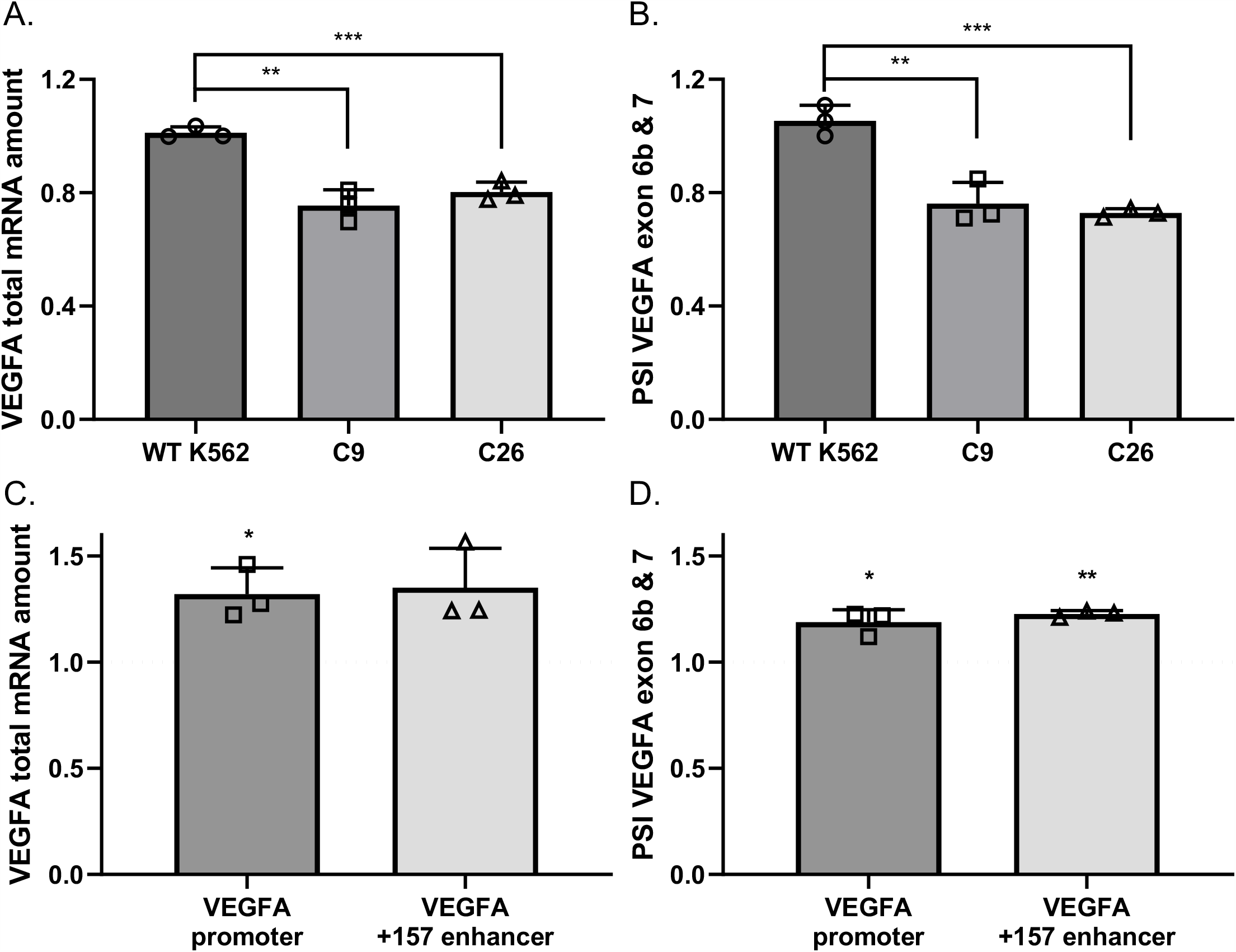
VEGFA +157 enhancer promotes inclusion of VEGFA exons 6b and 7. **A-B**. RNA was extracted from WT K562 cells and enhancer mutated clones 9 and 26, and analyzed by real-time PCR for total mRNA amount of VEGFA relative to *CycloA* and *hTBP* reference genes (**A**) and for VEGFA_121_ and VEGFA_189_ relative to VEGFA total mRNA amount. PSI was calculated by VEGFA_189_ relative to VEGFA_121_ (**B**). **C-D**. K562 cells were transfected with either dCas9-p300 core (mut) or dCas9-p300 core (WT) with four gRNAs targeted to the VEGFA promoter or +157 enhancer for 30 h. Total RNA was extracted and analyzed by real-time PCR for total mRNA of VEGFA relative to *CycloA* and *hTBP* reference genes (**C**) and for VEGFA_121_ and VEGFA_189_ relative to VEGFA total mRNA amount. PSI was calculated by VEGFA_189_ relative to VEGFA_121_ (**D**). Values are expressed as dCas9-p300 core (WT) relative to dCas9-p300 core (mut) and horizontal broken lines indicate no change between dCas9-p300 core (WT) relative to dCas9-p300 core (mut). Plots represents the mean of three independent experiments and ± SD (* p<0.05; ** p<0.01, ***p<0.001).

### Tethering p300 to the enhancer and promoter

To strengthen the link between enhancer activity and alternative splicing, we sought to activate the VEGFA +157 enhancer. We used the nuclease-deficient Cas9 (dCas9) conjugated to the p300 enzymatic core [33]. The CRISPR/dCas9 system allows us to tether a chromatin protein core domain to specific chromosome locations using a pool of four gRNAs. A pool of gRNAs was shown empirically to perform better than a single gRNA [33]. We co-transfected K562 cells with dCas9-p300 core (mut) or dCas9-p300 core (WT) and four gRNA plasmids targeting either the VEGFA promoter, or the +157 enhancer. Our results demonstrate that tethering p300’s enzymatic core to either the VEGFA promoter or the enhancer region up-regulates VEGFA total mRNA by 20-30% (**Fig. 2C**). This moderate change in expression may be due to a low dynamic range in K562 cells where the expression of VEGFA is very high and the chromatin at the promoter and enhancer is highly accessible. Accessibility of the chromatin can be demonstrated by acetylation of H3K27 and DHS sites at these regions (**Fig. 1B**). In addition to total mRNA amount, we measured the amount of the enhancer RNA (eRNA) generated by the +157 enhancer. We designed primers to the 3188 nt long eRNA (GH06J043925) (**Supplementary Fig. S1**) predicted in the +157 region and found similar levels of eRNA and also of VEGFA nascent mRNA (**Supplementary Fig. S2B&C**). Monitoring VEGFA alternative splicing following tethering of p300 to the promoter or enhancer showed inclusion of VEGFA exons 6b and 7 (**Fig. 2D**). These results are in line with our previous results, with the mutated +157 enhancer, suggesting that VEGFA promoter and +157 enhancer promote inclusion of exons 6b and 7 giving rise to VEGFA_189_ and a reduction in VEGFA_121_. We have previously showed that VEGFA_189_ induces endothelial cell migration [34], and thus our results might point to the VEGFA +157 enhancer as promoter of expression and pro-angiogenic isoforms.

### High expression of +157 eRNA and inclusion of exons 6b and 7 in CML patients

VEGFA protein is secreted from CML cells, which leads to high levels of VEGFA in the plasma of patients [14, 35]. We asked if overexpression of VEGFA in CML patients is accompanied by a change in alternative splicing. We extracted RNA from peripheral blood of twelve patients at different stages of their disease, represented by BCR-ABL fusion gene quantification (**Supplementary Table S1**). Although we could not detect significantly higher expression levels of VEGFA total mRNA at diagnosis, we identified significant overexpression of the +157 enhancer eRNA in a subset of patients, suggesting activation of the enhancer (**Fig. 3A)**. Specifically, our results show that both enhancer activation and inclusion of exons 6b and 7 are reduced when patients are in remission, suggesting that in CML patients, activity of the +157 enhancer goes hand in hand with a change in alternative splicing (**Fig. 3A and Supplementary Fig. S3**).

**Fig 3:**
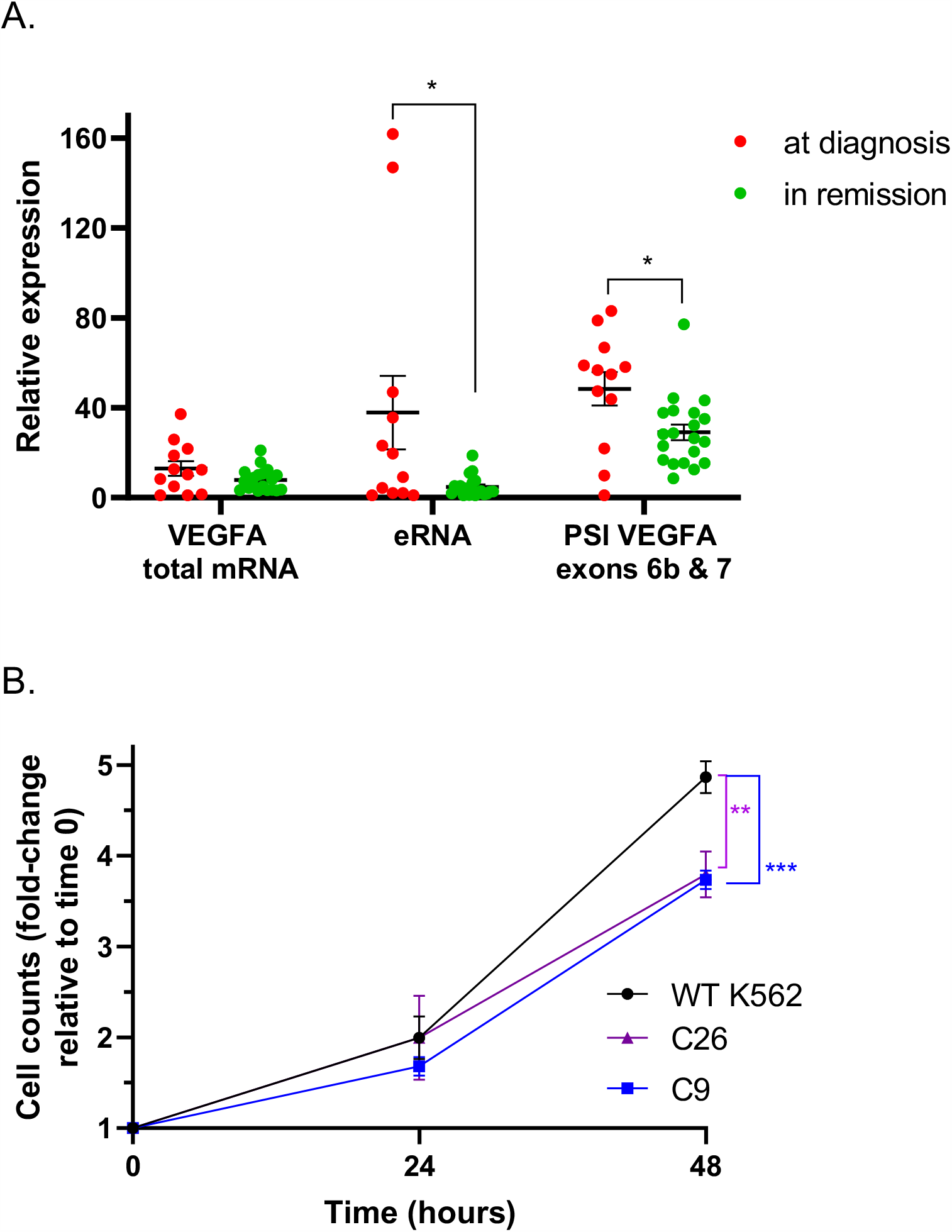
CML patients have greater inclusion of VEGFA exons 6b and 7 at diagnosis. **A**. RNA samples were collected from 12 CML patients at diagnosis (BCR-ABL positive, red circles) and during remission (BCR-ABL negative, green circles). RNA was analyzed by real time PCR for total mRNA amount of VEGFA and VEGFA +157 eRNA relative to *CycloA* and *hTBP* reference genes and for VEGFA_121_ and VEGFA_189_ relative to VEGFA total mRNA amount. PSI was calculated by VEGFA_189_ relative to VEGFA_121_. **B**. WT K562 cells and enhancer mutated C9 and C26 were seeded and counted every other day. Plot represents the mean of three independent experiments done with three replicates and ± SD (*p<0.05, Student’s t test).

### +157 enhancer is important for cell proliferation

We have previously shown that hypoxia and induction of VEGFA expression result in inclusion of exons 6b and 7 [34]. This suggests that the VEGFA_189_ isoform is important for physiological function and that its reduction following enhancer mutation could have an effect on cell proliferation. To demonstrate the physiological relevance of the enhancer-mediated alternative splicing of VEGFA, we monitored the proliferation rate of the parental K562 cells and the enhancer mutated clones 9 and 26. Our results demonstrate that mutations at VEGFA +157 enhancer in clones 9 and 26 resulted in slower proliferation rate (**Fig. 3B**). This suggests that the level of VEGFA expression as well as its alternative splicing is important for cell proliferation in a CML cell line.

### Slow RNAPII elongation rate promotes VEGFA exclusion of exons 6b and 7

Reduced expression of VEGFA as a result of enhancer +157 mutation could be due to less transcription initiation and/or slow RNAPII elongation kinetics along the gene. To examine the role of RNAPII elongation rate in the alternative splicing of VEGFA, we slowed RNAPII using a low concentration of CPT (6 µM) in K562 cells. Our results show that total VEGFA mRNA and eRNA were lower but no change in nascent mRNA was detected (**Fig. 4A and Supplementary Fig. S4A&B**). In addition, we found that slowing RNAPII elongation rate promotes exclusion of VEGFA exons 6b and 7 (**Fig. 4B**). Furthermore, we co-transfected HEK293T cells with plasmids expressing α-amanitin-resistant polymerases, a mutant “slow” RNAPII (mutant C4) or WT RNAPII [36]. α-amanitin blocks only endogenous RNAPII transcription as the WT and “slow” plasmids are resistant to it. We assessed the alternative splicing pattern of VEGFA generated by transcription with the recombinant polymerases. Our results show that total VEGFA mRNA and VEGFA nascent RNA were lower with no change in eRNA, and in addition we observed exclusion of VEGFA exons 6b and 7 (**Supplementary Fig. S4C-F**). These findings are similar to our findings with VEGFA +157 mutated enhancer (**Fig. 2B**). This suggests that the *VEGFA* enhancer promotes reduced RNAPII elongation rate to affect both gene expression and alternative splicing of the *VEGFA* gene.

**Fig 4:**
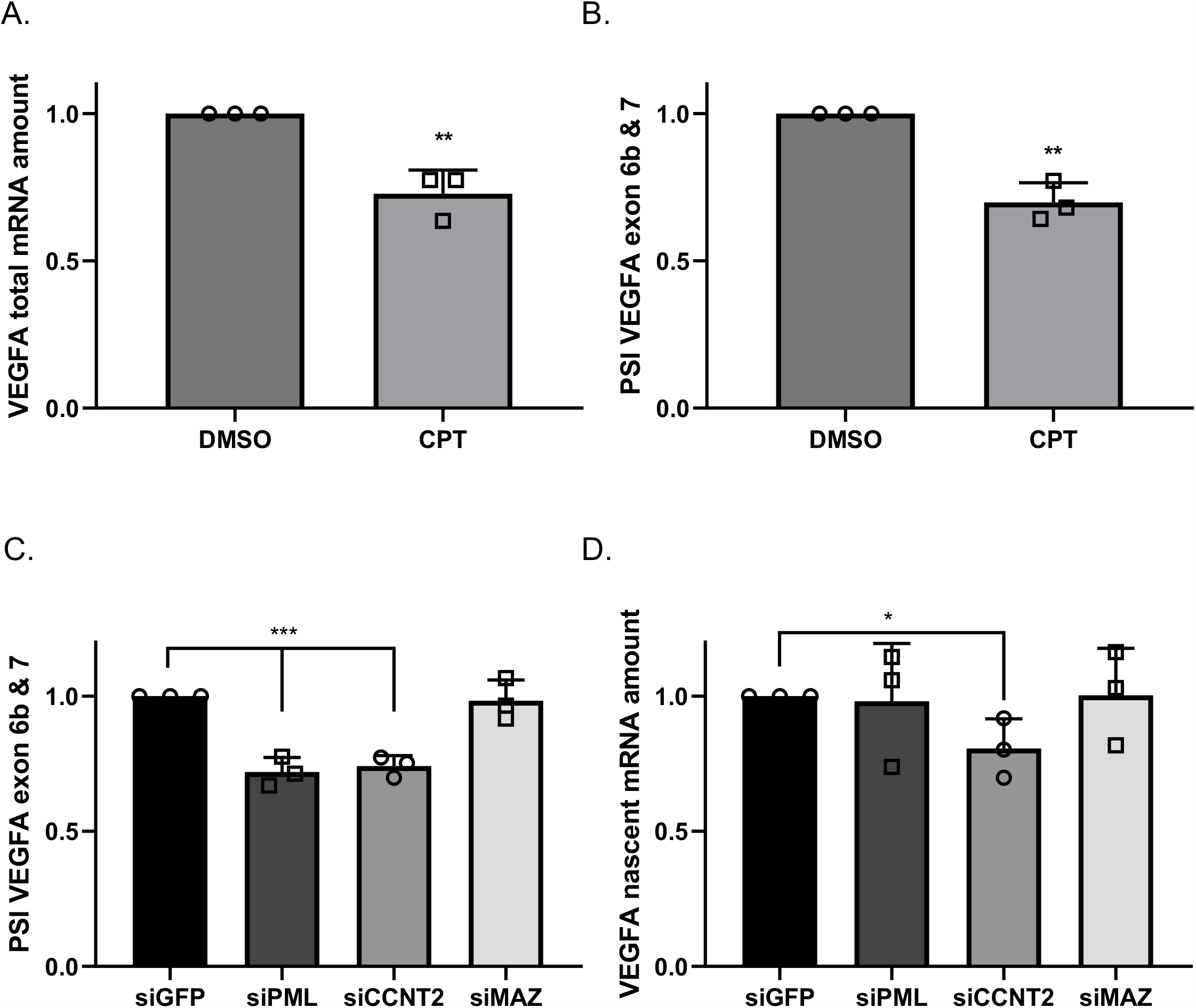
Slowing elongation of RNAPII and silencing its positive regulator CCNT2 promote exclusion of VEGFA exons 6b and 7. **A-B**. K562 cells were treated with 6 µM CPT for 6 h. Total RNA was extracted and analyzed by real-time PCR for total mRNA amount of VEGF relative to *CycloA* and *hTBP* reference genes (**A**) and for and for VEGFA_121_ and VEGFA_189_ relative to VEGFA total mRNA amount (**B**). PSI was calculated by VEGFA_189_ relative to VEGFA_121_. **C-D**. K562 cells were transfected with siRNA against GFP as negative control and siRNA targeting PML, CCNT2 and MAZ for 72 h. RNA was analyzed by real time PCR for VEGFA_121_ and VEGFA_189_ relative to VEGFA total mRNA amount. PSI was calculated by VEGFA_189_ relative to VEGFA_121_ (**C**); and for nascent pre-mRNA VEGFA relative to VEGFA total mRNA amount (**D**). Values represent mean ± SD of three independent experiments (* p<0.05; ** p<0.01, ***p<0.001).

**Fig 5:**
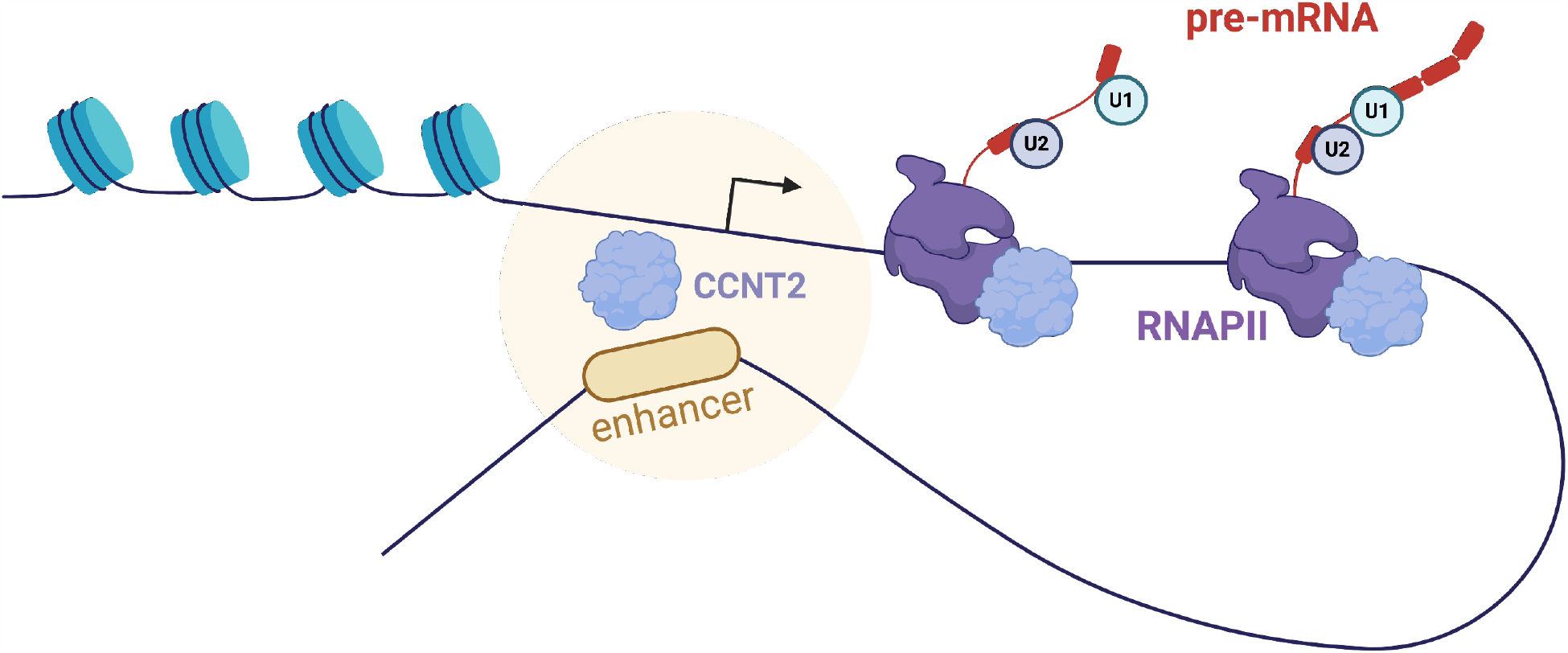
Schematic representation of the proposed mechanism. The RNAPII elongation facilitator CCNT2 binds to *VEGFA* promoter and enhancer and drive inclusion of exons 6b and 7. Spliceosome assembly on the nascent pre-mRNA is depicted by U1 and U2 short non-coding RNA.

### Chromatin factors role in VEGFA alternative splicing

Transcription factors are present in the spliceosome and are predicted to have a role in alternative splicing [37, 38]. To search for factors mediating the enhancer’s effect on alternative splicing, we analyzed 161 chromatin immunoprecipitation (ChIP)-seq experiments for DNA binding factors in K562 cells [39]. We set the cutoff of the signal binding normalized scores (in the range of 0-1000) to 350 and identified promyelocytic leukemia (PML), cyclin T2 (CCNT2) and MYC-associated zinc finger protein (MAZ) as binding to both the +157 enhancer and the promoter of the VEGFA gene (**Supplementary Table S2** and **Supplementary Fig. S4G&H**). PML co-localizes to PML nuclear bodies where it functions as a transcription factor and tumor suppressor [40]. CCNT2 is a cyclin that acts with its kinase partner CDK9 to regulate RNAPII transcription elongation [41, 42]. MAZ is a transcription factor with a known DNA binding motif and targets [43]. To check whether those proteins have a role in VEGFA alternative splicing, we silenced each of the factors in K562 cells with a pool of four siRNA oligos (**Supplementary Fig. S4I**). Our results show that no significant change was detected in VEGFA total mRNA amount but that PML and MAZ silencing caused a mild reduction in eRNA levels (**Supplementary Fig. S4J&K**). In addition, we detected exclusion of VEGFA exons 6b and 7 when silencing PML and CCNT2 but not when silencing MAZ (**Fig. 4C)**. Our results point to PML and CCNT2 as regulators of VEGFA alternative splicing, and their effect of exclusion of VEGFA exons 6b and 7, when silenced, is similar to that of mutating the +157 VEGFA enhancer (**Fig. 2B**). This similar effect on alternative splicing could suggest that the mutations in clones 9 and 26 are at the same location as PML and CCNT2 binding sites. Indeed, the mutation and the protein binding sites are found in the same DHS region (150 nt long) in close proximity (**Supplementary Fig. S4L**). CCNT2 is a positive regulator of RNAPII elongation kinetics [41, 42], and its silencing did yield a mild reduction in VEGFA nascent mRNA (**Fig. 4D**). Therefore silencing of CCNT2 leads to exclusion of VEGFA exons 6b and 7 likely by slowing down RNAPII elongation rate. Thus, our results suggest that the VEGFA enhancer regulate VEGFA splicing by regulating RNAPII elongation rate.

### Material and methods Cell lines and plasmids

HEK293T (ATCC Number: CRL-3216) and K562 (ATCC Number: CCL-243) cells were grown in Dulbecco’s modified Eagle’s medium (DMEM) and RPMI-1640, respectively, supplemented with 10% fetal bovine serum. Cell lines were maintained at 37°C and 5% CO_2_ atmosphere. Cells were transfected with TransIT-X2 Transfection Reagent (Roche) following the manufacturer’s instructions. After 30 h in culture, plasmid-transfected cells were used for experimentation. pcDNA-dCas9-p300 core plasmids (D1399Y; plasmid #61358 and plasmid #61357) were purchased from Addgene. pSPgRNA (Addgene, plasmid #47108) was used as the gRNA plasmid. The oligonucleotides containing the target sequences were hybridized, phosphorylated and cloned into the plasmid using BbsI sites. The target sequences are provided in Supplementary Table S3.

### RNAi

OnTarget Plus SMART pool of four siRNA oligomers per gene against PML, CCNT2 and MAZ were purchased from Sigma. Cells were grown to 20-50% confluence and transfected with siRNA using nucleofection (Amaxa). After 72 h in culture, siRNA transfected cells were used for experimentation.

### qRT-PCR

RNA was isolated from cells using the GENEzol TriRNA Pure Kit (GeneAid). cDNA synthesis was carried out with the Quanta cDNA Reverse Transcription Kit (QuantaBio). qPCR was performed with the iTaq Supermix (BioRad) on the BioRad iCycler. The collection of patient samples was approved by the institutional ethics committee of the Hadassah Medical Center. For *BCR-ABL* quantification, mononuclear cells derived from peripheral blood of CML patients were isolated on Ficoll-Hypaque gradient and total RNA was isolated. Real-time for *BCR-ABL* quantification was performed using ipsogen® BCR-ABL1 Mbcr IS-MMR (Qiagen). The comparative Ct method was employed to quantify transcripts, and delta Ct was measured in triplicate. Primers used in this study are provided in Supplementary Table S3.

### CPT treatment

To impede the dynamics of transcribing RNAPII, cells were treated with camptothecin (CPT, Sigma) to a final concentration of 6 µM for 6 h.

### Cell proliferation

Cell suspensions at a cell density of 2×10^4^/ml of K562 cells in logarithmic growth were seeded in 96-well cell culture plates. Cell cultures were divided into three groups with three triplicates for each group. The study group was incubated with sense and antisense nucleotides respectively, whereas the control group was incubated with RPMI-1640 at a final concentration of oligonucleotide of 4 µm. Cells were harvested at 24 h and 48 h after incubation. Cell viability was assessed using 0.4% trypan blue staining. The calculation was repeated for three times for each well to obtain a mean value.

## Discussion

Pre-mRNA splicing occurs co-transcriptionally [45], with the processes being coupled via kinetic and physical interaction of the RNA splicing machinery with RNAPII [38, 46]. Here we show that manipulating the VEGFA enhancer leads to regulation of its alternative splicing. We identify PML and CCNT2 as possible players mediating the connection of the enhancer’s activity with alternative splicing regulation. CCNT2 release pausing of RNAPII [41, 42] and indeed silencing it in K562 cells led to a mild reduction in VEGFA nascent RNA (**Fig. 4D**). Our results suggest that binding of CCNT2 to both VEGFA enhancer and promoter allow for fast elongation rate that promote high gene expression and inclusion of exons 6b and 7 (**Fig. 4B &C and Supplementary Fig. S4D**). Our results suggest that CCNT2 and PML is needed for VEGFA overexpression and thus might be upregulated in CML. While there is no data on CCNT2 in CML, it was shown to be overexpressed in acute myeloid leukemia and to promote proliferation in this cancer [47]. PML, while downregulated in many types of cancer, was found to be upregulated in CML and to be a positive regulator of self-renewal in CML-initiating cells [48]. Our work adds a level of complexity to alternative splicing regulation and deepens the interplay between transcription and alternative splicing.

VEGFA enhancer is demethylated in various hematopoietic cancers, resulting in higher gene expression [2]. Our work suggests that enhancer activation alters VEGFA alternative splicing to produce isoforms that promote angiogenesis. The idea of enhancers making a connection between gene expression and isoform abundance seems highly intuitive on the physiological level. This connection is strengthened by our observation that in CML patients we observed inclusion of exons 6b and 7 (**Fig. 3A**), which promotes VEGFA_189_ that we have previously shown to induce endothelial cell migration [34]. Finally, VEGFA +157 enhancer promotes VEGFA isoforms that have been shown to promote endothelial cell migration and thus cancer.

Our previous work connected promoter activity to alternative splicing by the acetyltransferase p300. Binding of p300 at the promoter region acetylates not only histones at this region, but also splicing factors so regulating alternative splicing of the specific gene [49]. Acetylation of splicing factors can either weaken or strengthen the splicing factor’s RNA binding properties that in turn regulate alternative splicing. Our results tethering p300 to the promoter and +157 enhancer of VEGFA in K562 cells show inclusion of exons 6b and 7 (**Fig. 2D**). This result attests to the activation of the enhancer and its effect on splicing also suggests that acetylating the histones at the promoter and enhancer regions allows stronger binding of transcription factors that then regulate alternative splicing, as we show for PML and CCNT2. Furthermore, this result could also indicate that p300 binding to the enhancer loops it to the promoter to allow for acetylation of splicing factors. Additional work is needed to connect these multiple layers of alternative splicing regulation.

Our work here focuses on one enhancer of the VEGFA gene, but chromatin marks and Hi-C data collected in K562 cells suggest that VEGFA has at least five more enhancers located in proximity to the +157 enhancer (**Fig. 1B**). Hence, the expression of *VEGFA* is the sum of inputs from its multiple enhancers. This could explain the mild change in expression we observed when tethering p300 to the +157 enhancer (**Fig. 2C**). Our hypothesis is that alternative splicing regulation by the enhancer is also the sum of several enhancers.

Our results show weak changes in eRNA while manipulating the enhancer. Activating the enhancer by tethering p300 did not show any change in eRNA expression (**Supplementary Fig. S2C**), but silencing PML transcription factor which binds VEGFA promoter and +157 enhancer show a mild reduction in eRNA and VEGFA exclusion of exons 6b and 7 (**Fig. 4C&D and Supplementary Fig. S4J&K**). These results do not form a strong association between the eRNA and alternative splicing and additional experiments are needed to further study this connection.

## Acknowledgments

We thank Dina Ben-Yehuda for valuable discussion and sharing her knowledge about the patients’ care and symptoms, and Svetelana Krichevsky for patient samples and for sharing her vast knowledge. We thank Asaf Hellman and Yehudit Bergman for the enhancer mutated K562 cells and advice throughout this work. We thank Alberto Kornblihtt for slow RNAPII plasmids.

---

**Fig S1:**
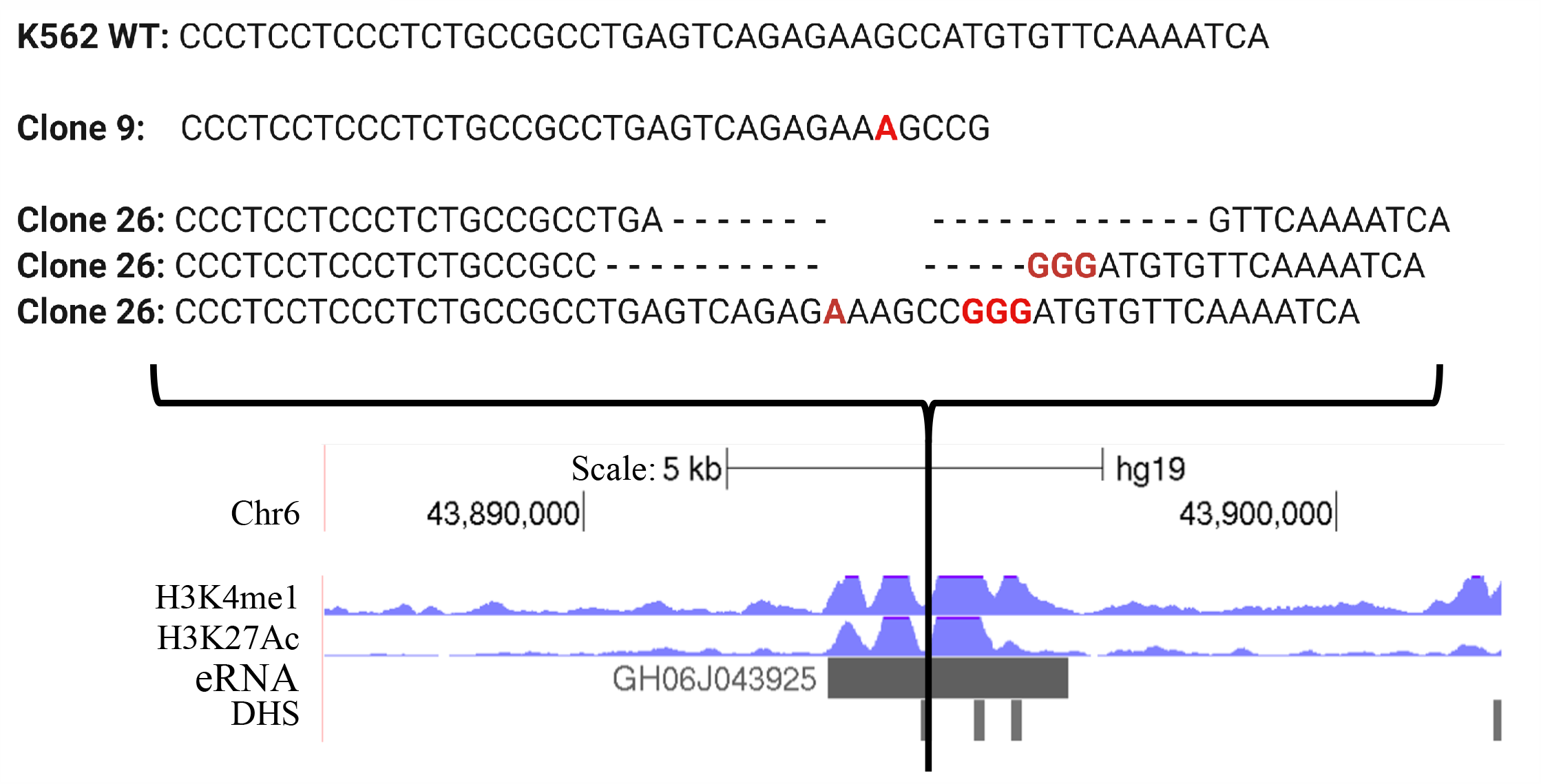
Sequence alignments of the VEGF +157 enhancer site showing K562 WT, C9 and C26 clones. A black line indicate the location of the mutations at the the VEGFA locus. ChIP-seq tracks for H3K27ac and H3K4me1 including DHS at the VEGFA locus in K562 cells. VEGFA +157 enhancer is marked by the eRNA (GH06J043925) and is enriched for H3K27ac and H3K4me1.

**Fig S2:**
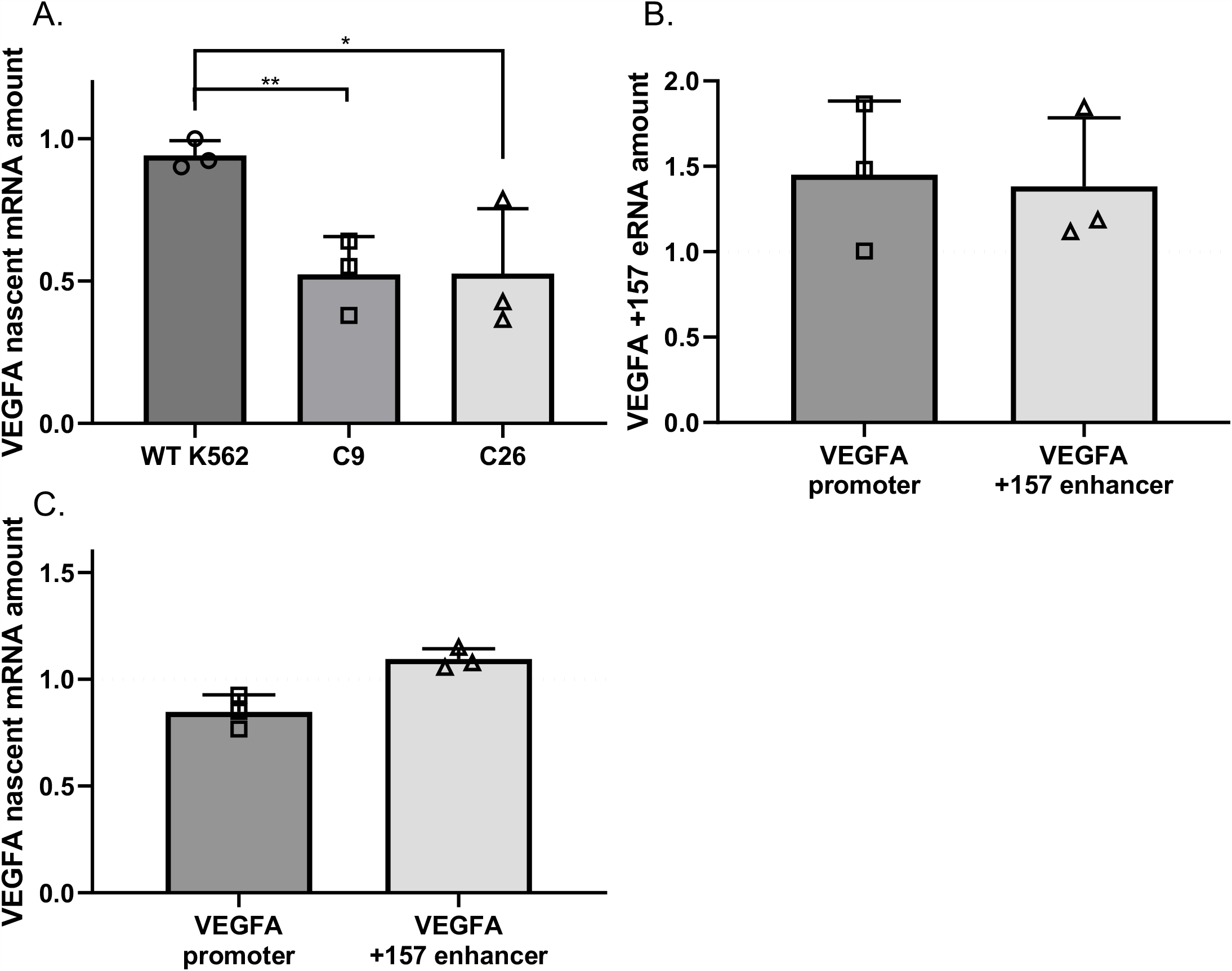
**A**. RNA was extracted from WT K562 cells and enhancer mutated C9 and C26, and analyzed by real-time PCR for nascent pre-mRNA VEGFA relative to VEGFA total mRNA amount. **B-C**. K562 cells were transfected with either dCas9-p300 core (mut) or dCas9-p300 core (WT) with four gRNAs targeted to the VEGFA promoter or +157 enhancer for 30 h. Total RNA was extracted and and analyzed by real-time PCR for total mRNA amount of for VEGFA +157 eRNA relative to *CycloA* and *hTBP* reference genes (**B**); and for nascent pre-mRNA VEGFA relative to VEGFA total mRNA amount (**C**); and horizontal broken lines indicate no change between dCas9-p300 core (WT) relative to dCas9-p300 core (mut). Values are expressed as dCas9-p300 core (WT) relative to dCas9-p300 core (mut). Plots represents the mean of three independent experiments and ±SD (* p<0.05; ** p<0.01).

**Fig S3:**
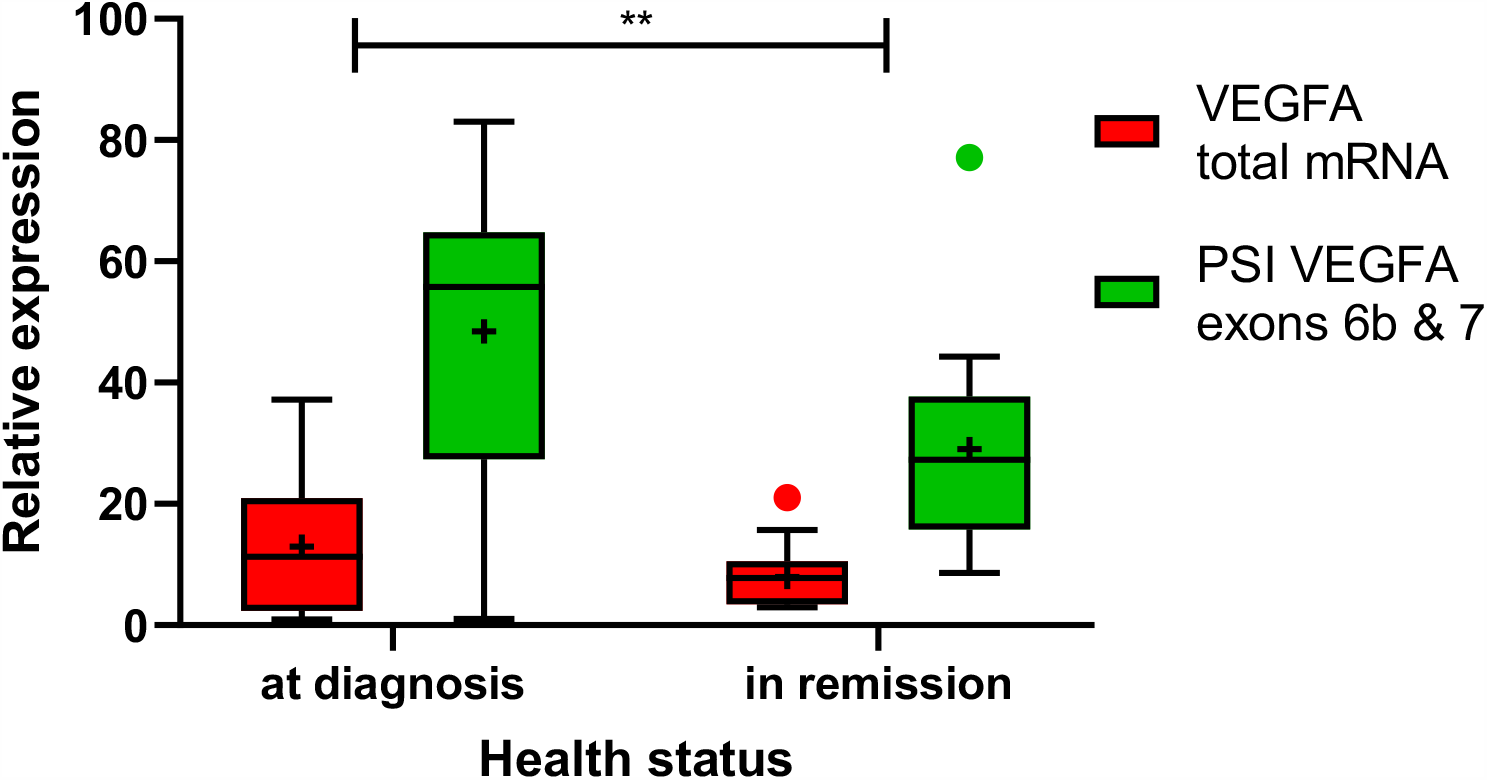
RNA samples were collected from 12 CML patients at diagnosis (BCR-ABL positive, diagnosis) and during remission (BCR-ABL negative, remission). RNA was analyzed by real time PCR for total mRNA amount of VEGFA relative to *CycloA* reference gene and for VEGFA_121_ and VEGFA_189_ relative to VEGFA total mRNA amount. PSI was calculated by VEGFA_189_ relative to VEGFA_121_. The results are represented as a Tukey boxplot. Analysis by mixed effects model (REML) showed a significant difference in relative expression between diagnosis and remission (predicted mean at diagnosis: 30.57; predicted mean in remission: 18.50; difference between predicted means: 12.07 ± 3.79, 95% CI of difference 4.21-19.93, p=0.0043).

**Fig S4:**
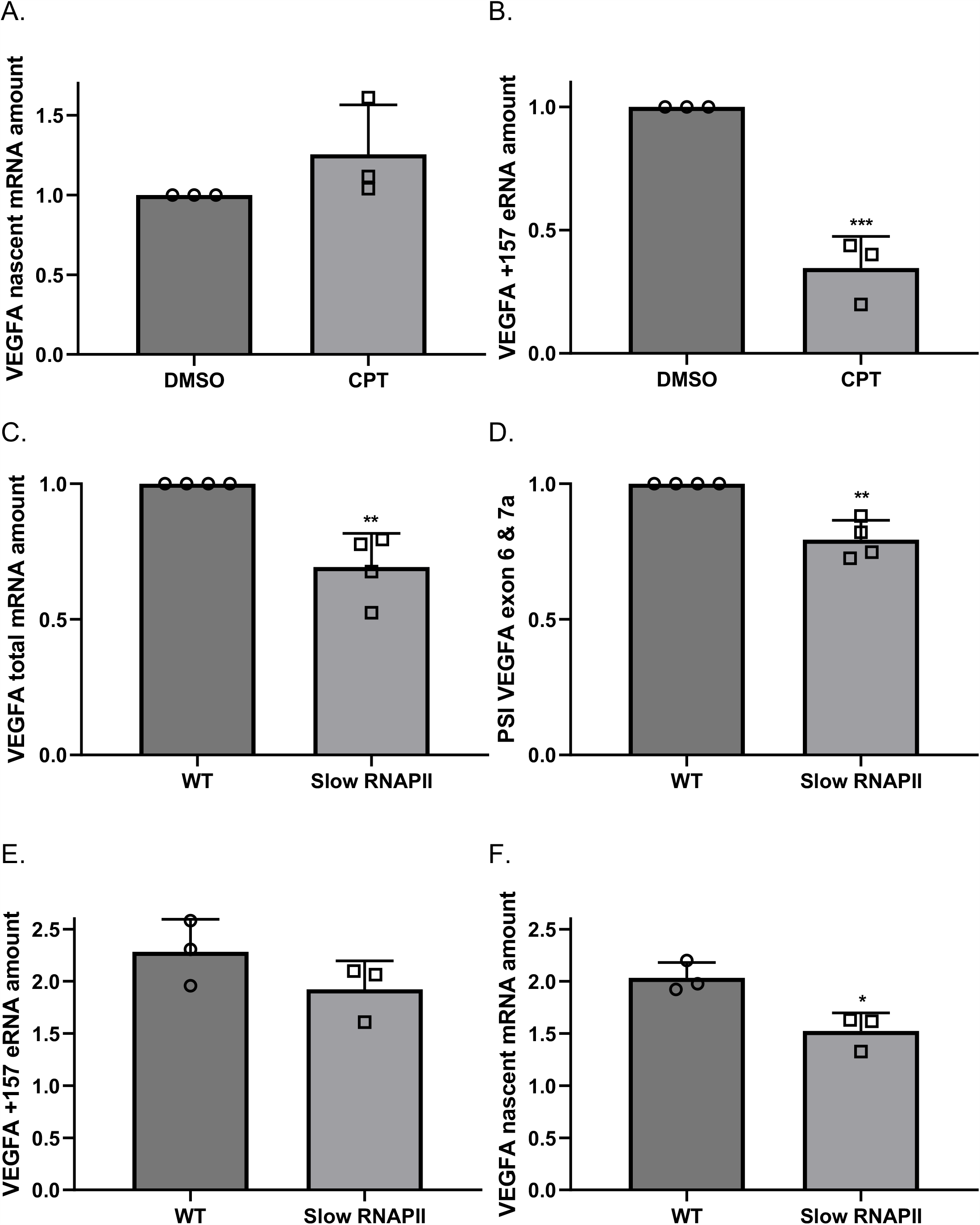
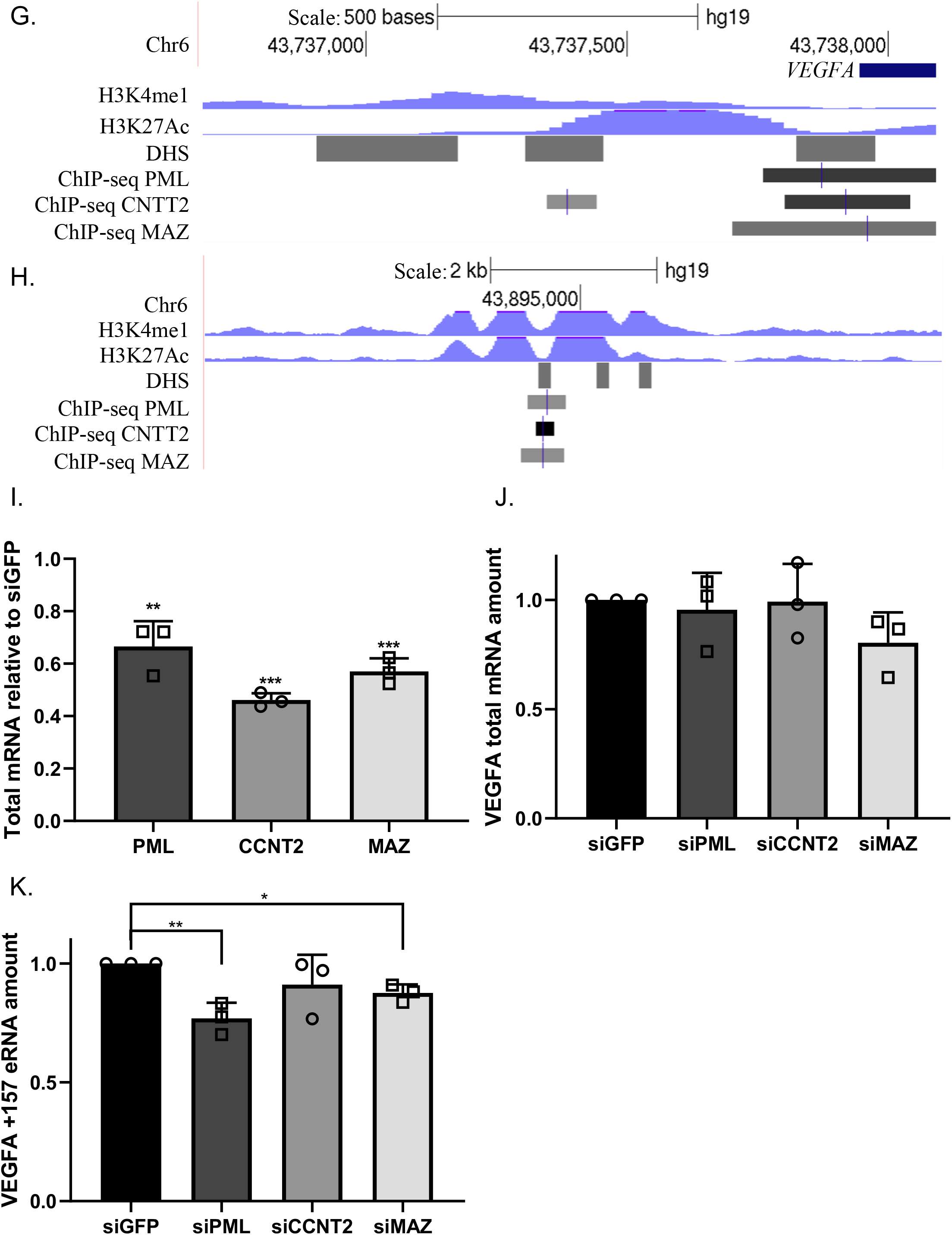
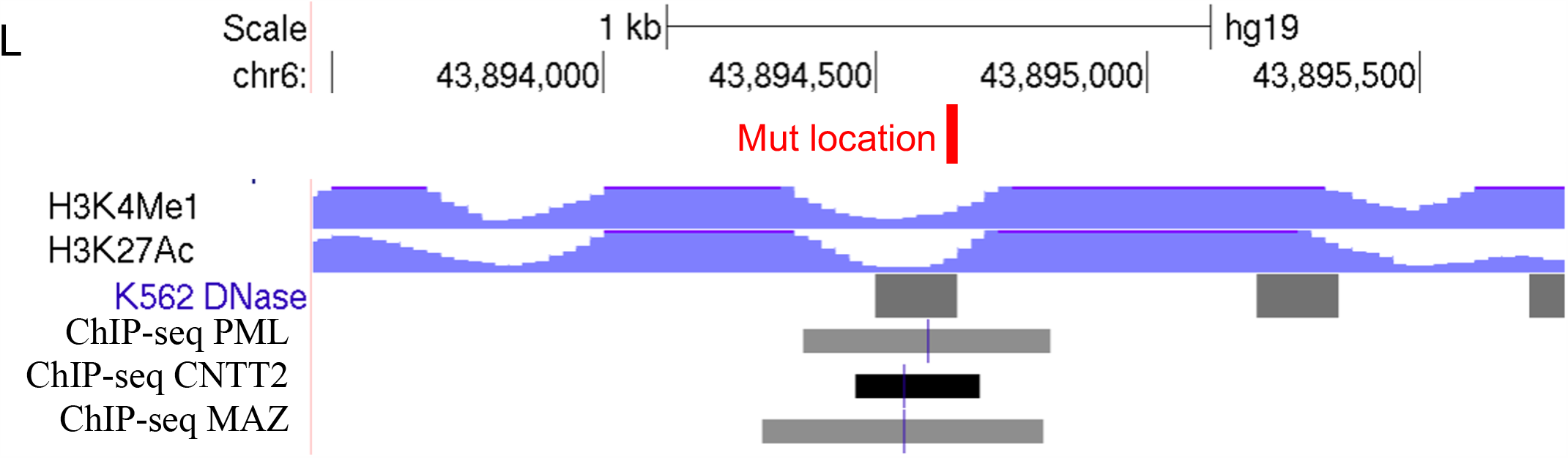
**A-D**. HEK293T cells were transfected with either WT RNAPII or slow RNAPII and after 24 h treated with α-amanitin for 24 h. Total RNA was extracted and analyzed by real-time PCR for total mRNA amount of VEGF relative to *CycloA* and *hTBP* reference genes (**A**) and VEGFA_121_ and VEGFA_189_ relative to VEGFA total mRNA amount. PSI was calculated by VEGFA_189_ relative to VEGFA_121_ (**B**) and for VEGFA +157 eRNA relative to *CycloA* and *hTBP* reference genes (**C**) and for nascent pre-mRNA VEGFA relative to VEGFA total mRNA amount (**D**). **E-F**. K562 cells were treated with 6 µM CPT for 6 h. Total RNA was extracted and analyzed by real-time PCR for VEGFA +157 eRNA relative to *CycloA* and *hTBP* reference genes (**E**) and for nascent pre-mRNA VEGFA relative to VEGFA total mRNA amount. **G-H**. ChIP-seq tracks for H3K27ac, H3K4me1, PML, CNTT2 and MAZ including DHS and at the VEGFA promoter (**G**) and VEGFA +157 enhancer (**H**) locus in K562 cells. **I-K**. K562 cells were transfected with siRNA against GFP as negative control and siRNA targeting PML, CNTT2 and MAZ for 72 h. RNA was analyzed by real time PCR for PML, CNTT2 and MAZ (**I**) and for VEGFA total mRNA amount relative to *CycloA* and *hTBP* reference genes (**J**) and for VEGFA +157 eRNA relative to *CycloA* and *hTBP* reference genes (**K**). Values represent mean ± SD of four (A-D) or three (E-F and I-K) independent experiments (* p<0.05; ** p<0.01, ***p<0.001).

**Supplementary Table S1:**
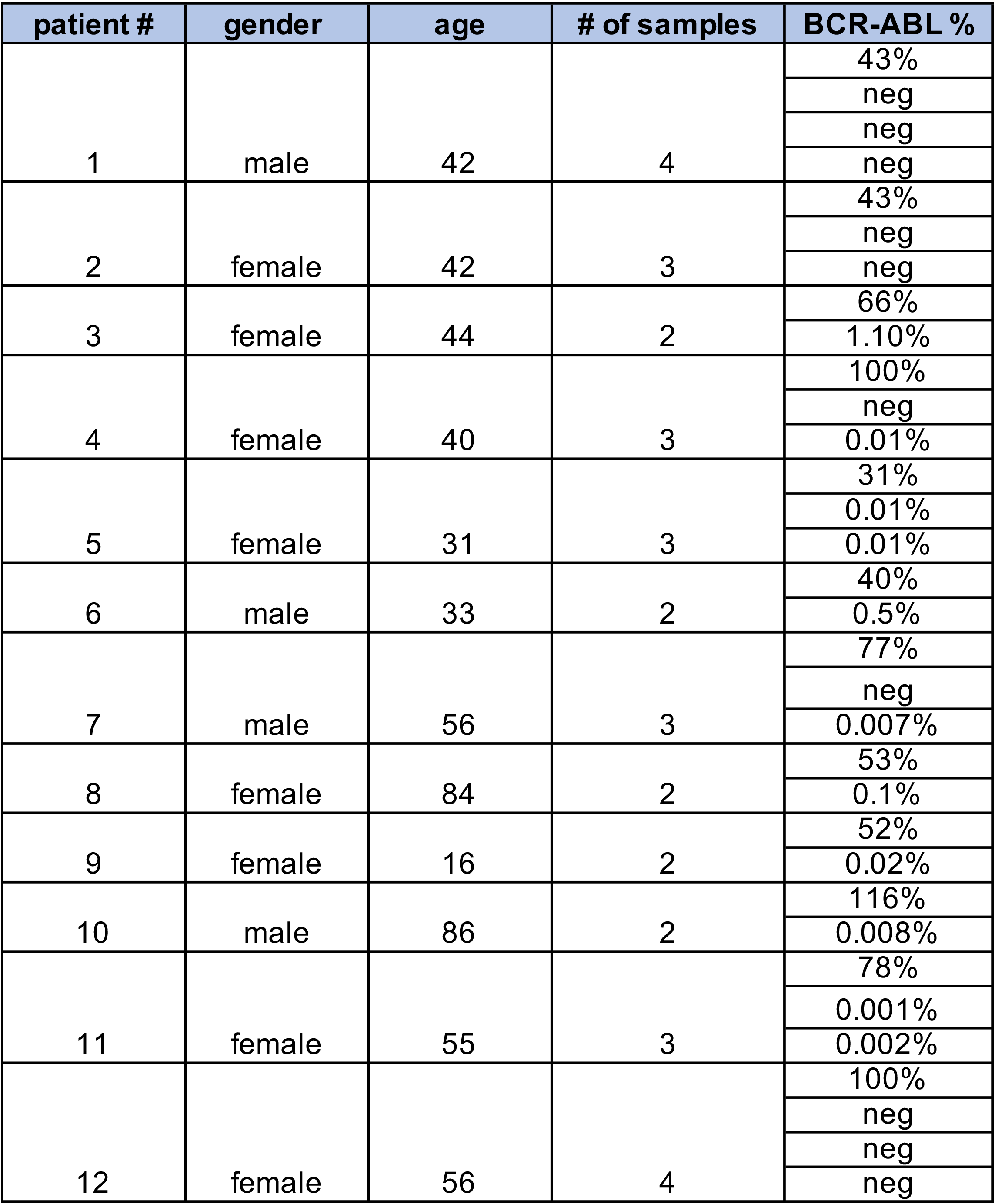
Details of patients and BCR-ABL %

**Supplementary Table S2:**
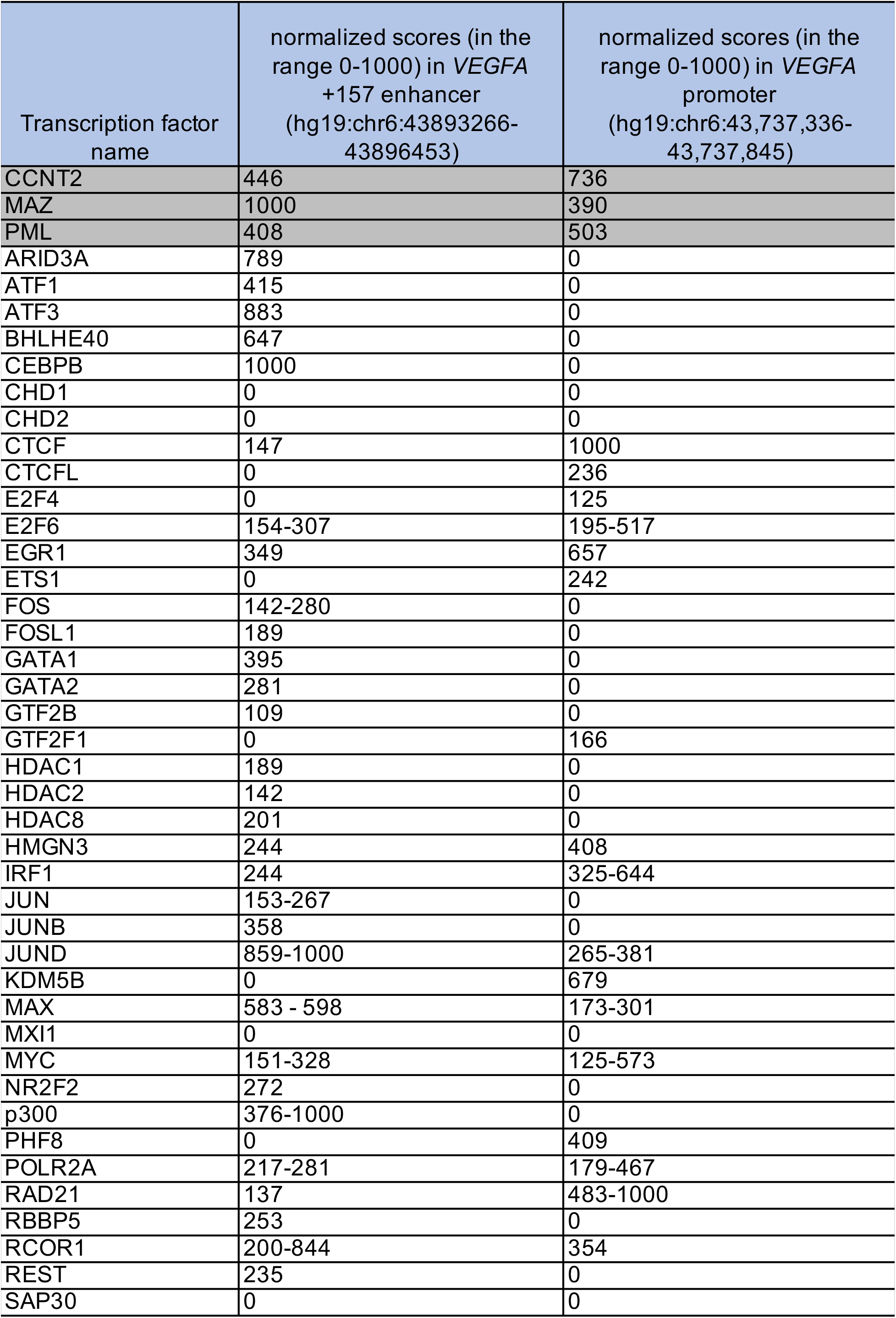
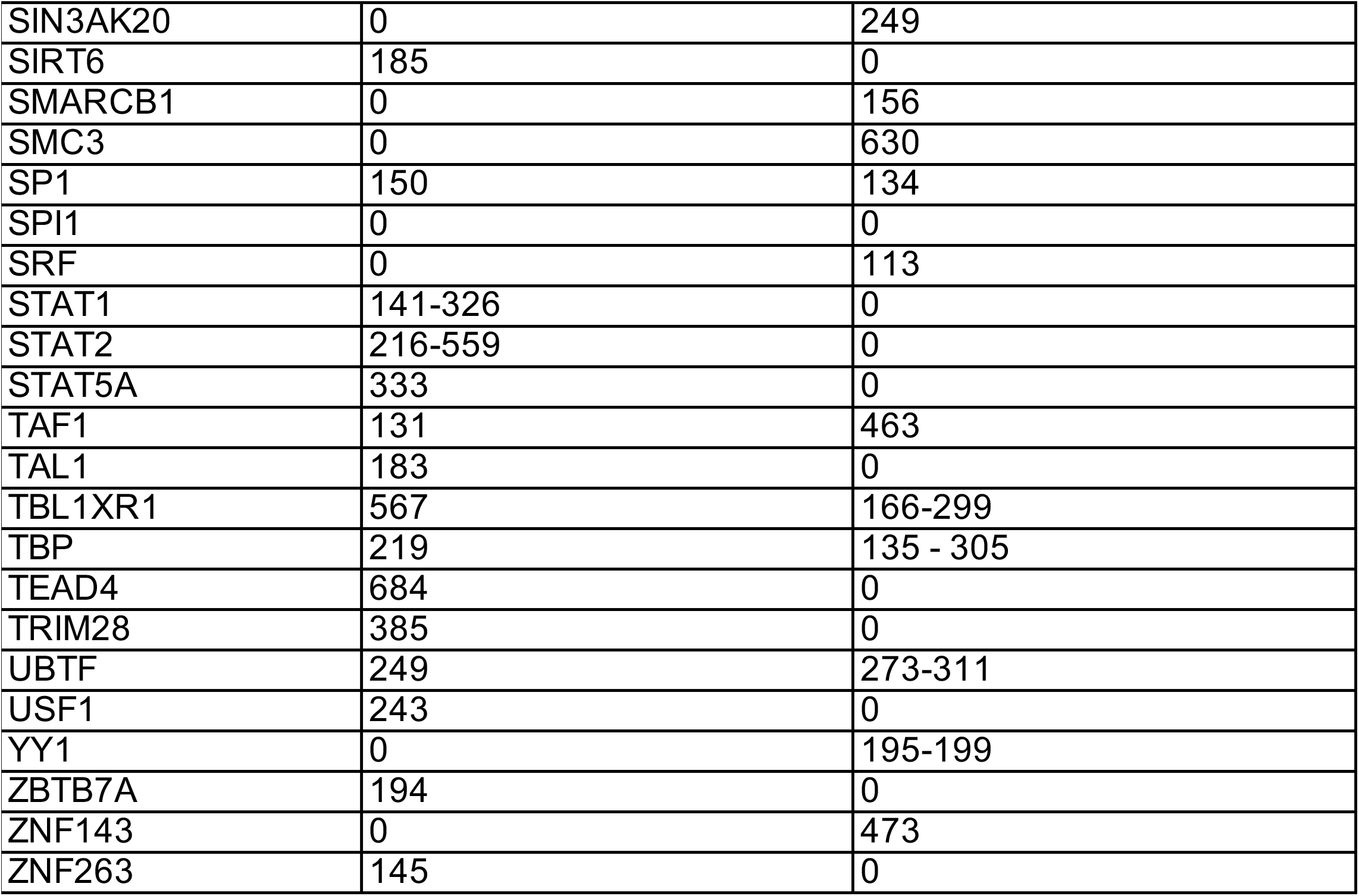
List of transcription and chromatin factors binding to *VEGFA* promoter and +157 enhancer

**Supplementary Table S3.**
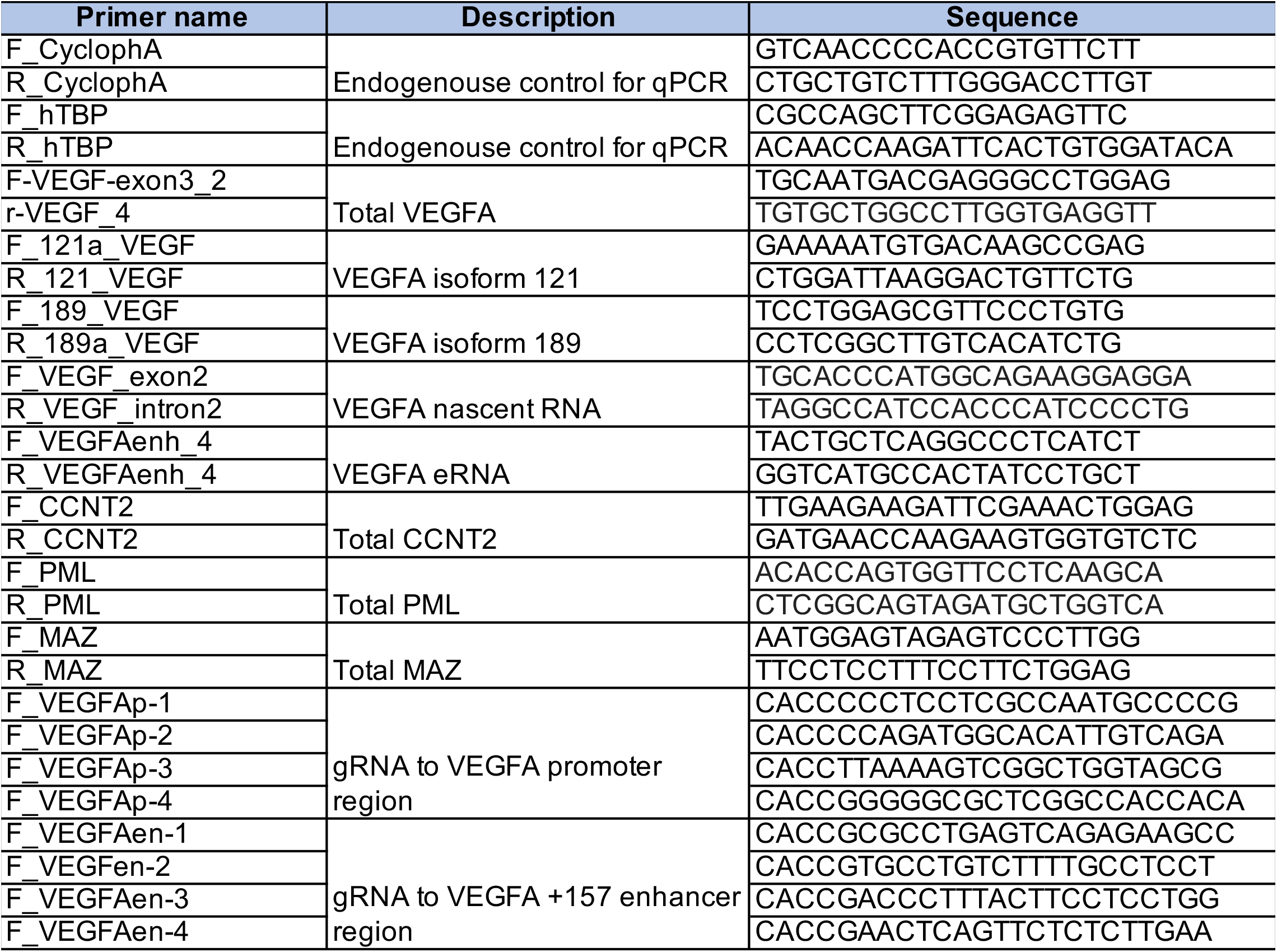
Primers used in this study

## References

1. Drier, Y., Enhancer and superenhancer regulation and its disruption in cancer. Current Opinion in Systems Biology, 2020. 19: p. 24–30.

2. Aran, D., et al., Embryonic Stem Cell (ES)-Specific Enhancers Specify the Expression Potential of ES Genes in Cancer. PLoS Genet, 2016. 12(2): p. e1005840.

3. Hicklin, D.J. and L.M. Ellis, Role of the vascular endothelial growth factor pathway in tumor growth and angiogenesis. J Clin Oncol, 2005. 23(5): p. 1011–27.

4. Nishida, N., et al., Angiogenesis in cancer. Vasc Health Risk Manag, 2006. 2(3): p. 213–9.

5. Ferrara, N., Vascular endothelial growth factor: basic science and clinical progress. Endocr Rev, 2004. 25(4): p. 581–611.

6. Folkman, J., How is blood vessel growth regulated in normal and neoplastic tissue? G.H.A. Clowes memorial Award lecture. Cancer Res, 1986. 46(2): p. 467–73.

7. Perez-Atayde, A.R., et al., Spectrum of tumor angiogenesis in the bone marrow of children with acute lymphoblastic leukemia. Am J Pathol, 1997. 150(3): p. 815–21.

8. Aguayo, A., et al., Angiogenesis in acute and chronic leukemias and myelodysplastic syndromes. Blood, 2000. 96(6): p. 2240–5.

9. Groffen, J., et al., Philadelphia chromosomal breakpoints are clustered within a limited region, bcr, on chromosome 22. Cell, 1984. 36(1): p. 93–9.

10. Hehlmann, R., et al., Assessment of imatinib as first-line treatment of chronic myeloid leukemia: 10-year survival results of the randomized CML study IV and impact of non-CML determinants. Leukemia, 2017. 31(11): p. 2398–2406.

11. Jabbour, E. and H. Kantarjian, Chronic myeloid leukemia: 2020 update on diagnosis, therapy and monitoring. Am J Hematol, 2020. 95(6): p. 691–709.

12. Mayerhofer, M., et al., BCR/ABL induces expression of vascular endothelial growth factor and its transcriptional activator, hypoxia inducible factor-1alpha, through a pathway involving phosphoinositide 3-kinase and the mammalian target of rapamycin. Blood, 2002. 100(10): p. 3767–75.

13. Janowska-Wieczorek, A., et al., Bcr-abl-positive cells secrete angiogenic factors including matrix metalloproteinases and stimulate angiogenesis in vivo in Matrigel implants. Leukemia, 2002. 16(6): p. 1160–6.

14. Legros, L., et al., Imatinib mesylate (STI571) decreases the vascular endothelial growth factor plasma concentration in patients with chronic myeloid leukemia. Blood, 2004. 104(2): p. 495–501.

15. Ebos, J.M., et al., Imatinib mesylate (STI-571) reduces Bcr-Abl-mediated vascular endothelial growth factor secretion in chronic myelogenous leukemia. Mol Cancer Res, 2002. 1(2): p. 89–95.

16. Lakkireddy, S., et al., Association of Vascular Endothelial Growth Factor A (VEGFA) and its Receptor (VEGFR2) Gene Polymorphisms with Risk of Chronic Myeloid Leukemia and Influence on Clinical Outcome. Mol Diagn Ther, 2016. 20(1): p. 33–44.

17. Lundberg, L.G., et al., Bone marrow in polycythemia vera, chronic myelocytic leukemia, and myelofibrosis has an increased vascularity. Am J Pathol, 2000. 157(1): p. 15–9.

18. Wang, E. and I. Aifantis, RNA Splicing and Cancer. Trends Cancer, 2020. 6(8): p. 631–644.

19. Oltean, S. and D.O. Bates, Hallmarks of alternative splicing in cancer. Oncogene, 2014. 33(46): p. 5311–8.

20. Salton, M., et al., Inhibition of vemurafenib-resistant melanoma by interference with pre-mRNA splicing. Nat Commun, 2015. 6: p. 7103.

21. Salton, M. and T. Misteli, Small Molecule Modulators of Pre-mRNA Splicing in Cancer Therapy. Trends Mol Med, 2016. 22(1): p. 28–37.

22. Cramer, P., et al., Functional association between promoter structure and transcript alternative splicing. Proc Natl Acad Sci U S A, 1997. 94(21): p. 11456–60.

23. Roberts, G.C., et al., Co-transcriptional commitment to alternative splice site selection. Nucleic Acids Res, 1998. 26(24): p. 5568–72.

24. Kadener, S., et al., Regulation of alternative splicing by a transcriptional enhancer through RNA pol II elongation. Proc Natl Acad Sci U S A, 2002. 99(12): p. 8185–90.

25. Naftelberg, S., et al., Regulation of alternative splicing through coupling with transcription and chromatin structure. Annu Rev Biochem, 2015. 84: p. 165–98.

26. Dujardin, G., et al., How slow RNA polymerase II elongation favors alternative exon skipping. Mol Cell, 2014. 54(4): p. 683–90.

27. Carmeliet, P., et al., Impaired myocardial angiogenesis and ischemic cardiomyopathy in mice lacking the vascular endothelial growth factor isoforms VEGF164 and VEGF188. Nat Med, 1999. 5(5): p. 495–502.

28. Zhang, H.T., et al., The 121 amino acid isoform of vascular endothelial growth factor is more strongly tumorigenic than other splice variants in vivo. Br J Cancer, 2000. 83(1): p. 63–8.

29. Herve, M.A., et al., Overexpression of vascular endothelial growth factor 189 in breast cancer cells leads to delayed tumor uptake with dilated intratumoral vessels. Am J Pathol, 2008. 172(1): p. 167–78.

30. Grunstein, J., et al., Isoforms of vascular endothelial growth factor act in a coordinate fashion To recruit and expand tumor vasculature. Mol Cell Biol, 2000. 20(19): p. 7282–91.

31. Consortium, E.P., An integrated encyclopedia of DNA elements in the human genome. Nature, 2012. 489(7414): p. 57–74.

32. Vian, L., et al., The Energetics and Physiological Impact of Cohesin Extrusion. Cell, 2018. 175(1): p. 292–294.

33. Hilton, I.B., et al., Epigenome editing by a CRISPR-Cas9-based acetyltransferase activates genes from promoters and enhancers. Nat Biotechnol, 2015. 33(5): p. 510–7.

34. Salton, M., T.C. Voss, and T. Misteli, Identification by high-throughput imaging of the histone methyltransferase EHMT2 as an epigenetic regulator of VEGFA alternative splicing. Nucleic Acids Res, 2014. 42(22): p. 13662–73.

35. Chen, H., et al., Expression of VEGF and its effect on cell proliferation in patients with chronic myeloid leukemia. Eur Rev Med Pharmacol Sci, 2015. 19(19): p. 3569–73.

36. de la Mata, M., et al., A slow RNA polymerase II affects alternative splicing in vivo. Mol Cell, 2003. 12(2): p. 525–32.

37. Lai, M.C., B.H. Teh, and W.Y. Tarn, A human papillomavirus E2 transcriptional activator. The interactions with cellular splicing factors and potential function in pre-mRNA processing. J Biol Chem, 1999. 274(17): p. 11832–41.

38. Kornblihtt, A.R., et al., Multiple links between transcription and splicing. RNA, 2004. 10(10): p. 1489–98.

39. Consortium, E.P., A user’s guide to the encyclopedia of DNA elements (ENCODE). PLoS Biol, 2011. 9(4): p. e1001046.

40. Gamell, C., et al., PML tumour suppression and beyond: therapeutic implications. FEBS Lett, 2014. 588(16): p. 2653–62.

41. Napolitano, G., B. Majello, and L. Lania, Catalytic activity of Cdk9 is required for nuclear co-localization of the Cdk9/cyclin T1 (P-TEFb) complex. J Cell Physiol, 2003. 197(1): p. 1–7.

42. Baumli, S., et al., The structure of P-TEFb (CDK9/cyclin T1), its complex with flavopiridol and regulation by phosphorylation. EMBO J, 2008. 27(13): p. 1907–18.

43. Song, J., et al., Transcriptional regulation by zinc-finger proteins Sp1 and MAZ involves interactions with the same cis-elements. Int J Mol Med, 2003. 11(5): p. 547–53.

44. Arcondeguy, T., et al., VEGF-A mRNA processing, stability and translation: a paradigm for intricate regulation of gene expression at the post-transcriptional level. Nucleic Acids Res, 2013. 41(17): p. 7997–8010.

45. Brody, Y. and Y. Shav-Tal, Transcription and splicing: when the twain meet. Transcription, 2011. 2(5): p. 216–20.

46. Maniatis, T. and R. Reed, An extensive network of coupling among gene expression machines. Nature, 2002. 416(6880): p. 499–506.

47. Ke, S., et al., MicroRNA-192 regulates cell proliferation and cell cycle transition in acute myeloid leukemia via interaction with CCNT2. Int J Hematol, 2017. 106(2): p. 258–265.

48. Ito, K., et al., PML targeting eradicates quiescent leukaemia-initiating cells. Nature, 2008. 453(7198): p. 1072–8.

49. Siam, A., et al., Regulation of alternative splicing by p300-mediated acetylation of splicing factors. RNA, 2019. 25(7): p. 813–824.

